# The role of chromosomal inversions in speciation

**DOI:** 10.1101/211771

**Authors:** Z.L. Fuller, C.J. Leonard, R.E. Young, S.W. Schaeffer, N Phadnis

## Abstract

The chromosomal inversions of *D. persimilis* and *D. pseudoobscura* have deeply influenced our understanding of the evolutionary forces that shape natural variation, speciation, and selfish chromosome dynamics. Here, we perform a comprehensive reconstruction of the evolutionary histories of the chromosomal inversions in these species. We provide a solution to the puzzling origins of the selfish *Sex-Ratio* chromosome in *D. persimilis* and show that this *Sex-Ratio* chromosome directly descends from an ancestrally-arranged chromosome. Our results further show that all fixed inversions between *D. persimilis* and *D. pseudoobscura* were segregating in the ancestral population long before speciation, and that the genes contributing to reproductive barriers between these species must have evolved within them afterwards. We propose a new model for the role of chromosomal inversions in speciation and suggest that higher levels of divergence and an association with hybrid incompatibilities are emergent properties of ancestrally segregating inversions. These findings force a reconsideration of the role of chromosomal inversions in speciation, not as protectors of existing hybrid incompatibility alleles, but as fertile grounds for their formation.

---

Chromosomal inversions are rearrangements of segments of DNA, where the linear order of a group of genes is reversed with respect to the original order. In crosses between two species that differ by one or more chromosomal inversions, the resulting hybrids can experience meiotic chromosome pairing problems due to these inversions and, therefore, become sterile. Chromosomal inversions can, thus, potentially play an important role in the evolution in intrinsic postzygotic barriers between species. Understanding the extent to which such chromosomal rearrangements play a role in speciation is a longstanding and fundamental problem in evolutionary genetics (1–3).

A key experimental test of the role of chromosomal inversions in hybrid sterility is the chromosome-doubling test (1, 4). According to this test, if diploid hybrids that are heterozygous for chromosomal inversions suffer sterility due to chromosome pairing problems, then doubling the chromosome number of these individuals should provide a collinear partner for each chromosome and restore meiotic chromosome pairing and hybrid fertility. This test for the role of chromosomal inversion in hybrid sterility, which manifests as the conversion of sterile diploid hybrids into fertile tetraploid hybrids, has been successfully carried out many times in plants in laboratories and in nature (1, 5, 6). This widespread success of the chromosomal doubling test has cemented the role of chromosomal rearrangements in the evolution of reproductive isolation in plants.

In contrast, the chromosome-doubling test has been carried out only once in animals, in F1 hybrid males between *Drosophila persimilis* and its closest sister species *D. pseudoobscura*. *D. persimilis* and *D. pseudoobscura* differ by three fixed chromosomal inversions, and hybrid F1 males between these species are sterile. Spermatocytes in these hybrid F1 males experience chromosome-pairing problems and degenerate after the first meiotic division resulting in hybrid male sterility. Groups of tetraploid spermatocytes, however, are frequently observed in these hybrid F1 males. In these tetraploid spermatocytes, every chromosome can now potentially pair properly with a homologous chromosome, which should restore fertility in the male hybrids. The chromosomal doubling test in this case, however, shows no rescue of hybrid fertility. Chromosomes in tetraploid spermatocytes in *D. persimilis*-*D. pseudoobscura* hybrid F1 males experience pairing problems at the same rates as in diploid spermatocytes, and undergo the same degenerative process after meiotic divisions (4). This singular failure of the chromosome-doubling test provided a key experimental line of evidence that chromosomal inversions are not likely to play a direct role in speciation in animals (2).

Further experiments to understand the genetic architecture of hybrid sterility loci in *D. persimilis*-*D. pseudoobscura* hybrids revealed the importance of deleterious epistatic interactions between unlinked loci as the cause of hybrid sterility (7, 8). These results led to the development of the Dobzhansky-Muller model for the evolution of genic hybrid incompatibilities, which has garnered substantial empirical support and now forms the bedrock of our understanding of the evolution of intrinsic postyzgotic isolating mechanisms in speciation. Together with other theoretical arguments, this body of work involving *D. persimilis* and *D. pseudoobscura* firmly established the idea that genic incompatibilities, rather than chromosomal inversions, are primarily responsible for the evolution of hybrid sterility in animals. The reasons why the role of chromosomal inversions in speciation differs between plants and animals has been the subject of much speculation, and involves differences between plants and animals in aspects of self-fertilization, open developmental plans, and the presence of degenerate sex chromosomes (1).

Interestingly, recent studies in *D. persimilis* and *D. pseudoobscura*–the same hybridization that led to the demise of the role of chromosomal inversions in animal speciation–have led to the dramatic resurgence of a modified version of this idea. Two new empirical observations regarding the patterns of reproductive isolation and genetic divergence in *D. persimilis* and *D. pseudoobscura* are key to these developments: i) nearly all genes that contribute to reproductive isolation between these species are located among the fixed chromosomal inversion differences, and ii) the fixed chromosomal inversions between these species display higher genetic divergence than collinear regions of the genome (9–14). Under the modified versions of the chromosomal theory of speciation that attempt to explain these two patterns, genic incompatibilities still play a major role in the evolution of hybrid sterility, but chromosomal inversions facilitate this process indirectly by linking sets of genes together that contribute to reproductive isolation. These new theories shifted the focus from chromosome pairing problems to the recombination suppressive properties of chromosomal inversions in the context of animal speciation (15).

According to the first version put forth by Noor (11, 16), hybrid incompatibility genes may initially evolve more or less uniformly across collinear and inverted regions in the genomes of isolated populations. If these populations later re-hybridize on secondary contact, then any incompatible alleles will be selected against because they exact a fitness cost in the form of unfit hybrid progeny. Inversions, however, suppress recombination and can generate a large block of tightly linked loci. If an incompatible allele is associated with an inversion that carries other beneficial alleles, then it may be preserved in the face of gene flow. In contrast, any incompatible alleles contained within collinear regions are unlikely to be tightly linked to beneficial alleles that may help preserve them, and will be replaced by the compatible allele. After secondary contact and gene flow, the only hybrid incompatibility loci that persist will be those that are associated with chromosomal inversions. In addition, these inversions will appear more diverged than collinear regions because the latter will have been homogenized by pervasive gene flow after speciation. By invoking gene flow during secondary contact after speciation, this model can explain both the association of hybrid incompatibility genes with chromosomal inversions, and the higher genetic divergence displayed by chromosomal inversions in comparison to collinear regions as observed between *D. persimilis* and *D. pseudoobscura*. Another version of this idea proposed by Rieseberg (16) posits that chromosomal inversions may link together many small effect hybrid sterility alleles, which may add all of these small effect alleles together to cause substantial sterility in hybrids. Both models both invoke gene flow during secondary contact after speciation, and we refer to these sets of models together as the ‘gene flow after speciation’ models.

In contrast to the ‘gene flow after speciation’ models, which rely on gene exchange after speciation to explain the above patterns, a theoretical argument put forth by Navarro and Barton invokes gene flow during speciation (henceforth referred to as the ‘gene flow during speciation’ model) (17). The ‘gene flow during speciation’ model, however, considers a scenario where an incompatible allele is located within a chromosomal inversion that also carries alleles that are beneficial in one population but not the other. Normally, due to the cost of producing unfit hybrids, an incompatible allele is not expected to increase in frequency within populations connected by migration. According to this model, however, the fitness cost incurred by an incompatible allele due to producing unfit hybrids can be offset by the fitness advantage conferred by its linkage to beneficial alleles. Chromosomal inversions that carry incompatible alleles along with other alleles that are beneficial in one population but not the other may persist for a long time or even go to fixation within populations despite gene flow during speciation. The collinear regions continue to be homogenized by gene flow between the two populations during this time, and can lead to the association of hybrid incompatibility alleles with inversions, and the higher divergence of inversions relative to collinear regions. Together, the plausibility of the models based on gene flow during or after speciation in explaining these empirical patterns through a combination of gene flow and recombination suppression have led to the widespread acceptance of this revised view of the indirect contribution of chromosomal inversions in speciation.

Here, we dissect the evolutionary history of the chromosomal inversions in *D. persimilis* and *D. pseudoobscura* to show that all fixed chromosomal inversions between these species segregated in their ancestral population, and predated the divergence between these species by a remarkable length of time. These results are contrary to the current views on the origins of these inversions, and have important implications for the role of chromosomal inversions in speciation. In particular, our results suggest that it is unnecessary to invoke gene flow during or after speciation as necessitated by other models to explain the patterns of hybrid incompatibilities and divergence between these species. Our key insights into deciphering the evolutionary histories of these chromosomal inversions, however, came not from studying the fixed inversion differences between these species, but from resolving the origins of the surprising arrangement of the *Sex-Ratio* chromosome of *D. persimilis*. We, therefore, first explain our resolution to the evolutionary history of this *Sex-Ratio* chromosome, and then proceed to reconstruct the evolutionary history of other chromosomal inversions between *D. persimilis* and *D. pseudoobscura*.

*Sex-Ratio* chromosomes are variants of *X*-chromosomes that are often found at high frequencies within natural populations (18). Males that carry a *Sex-Ratio* chromosome eliminate nearly all *Y*-bearing sperm (19), and produce nearly all female offspring (*i.e.,* heavily distorted progeny *sex-ratios*). By distorting the balance of segregation in their favor in excess of Mendelian expectations, these *Sex-Ratio* can rapidly spread through populations even if they reduce the fitness of the individuals that carry them (20, 21). In the absence of opposing forces such as the evolution of suppressor alleles, these selfish *X*-chromosomes may even drive populations to extinction (22). Such *Sex-Ratio* (*SR*) chromosomes have been identified in many Dipteran species, and are almost always associated with one or more characteristic chromosomal inversions relative to the wild type, or *Standard* (*ST*) chromosomes (21).

When a new chromosomal inversion generates tight linkage between a segregation distorter allele and other alleles that enhance distortion (or alleles that neutralize suppressors-of-distortion), this produces a stronger driving chromosome that can supplant its weaker versions (23). This process sets up an expected order for the evolution of *Sex-Ratio* chromosomes: distorter alleles arise first, enhancers of distortion appear next, and chromosomal inversions that tie these together arrive last. This framework explains why most *Sex-Ratio* chromosomes are associated with derived inversions. Consistent with this expectation, the *D. persimilis SR* chromosome is inverted with respect to the *D. persimilis ST* chromosome on the right arm of the *X* chromosome (*XR*). However, the *Standard D. persimilis XR* differs from *D. pseudoobscura XR* by a single derived inversion. Surprisingly, the *D. persimilis SR* inversion appears to have reversed the same derived *D. persimilis ST* inversion, such that *D. persimilis SR* appears collinear with *D. pseudoobscura* (Figure 1A). This unexpected collinearity of the *D. persimilis SR* chromosome with the *Standard* chromosome of its sister species is thought to be the result of a second inversion event on the background of *D. persimilis ST* at approximately the same breakpoints as the original *D. persimilis XR* inversion. However, previous molecular evolutionary studies have yielded conflicting results, and the origin of the *D. persimilis Sex-Ratio* inversion remains the subject of speculation (9, 24, 25).

**Figure 1:**
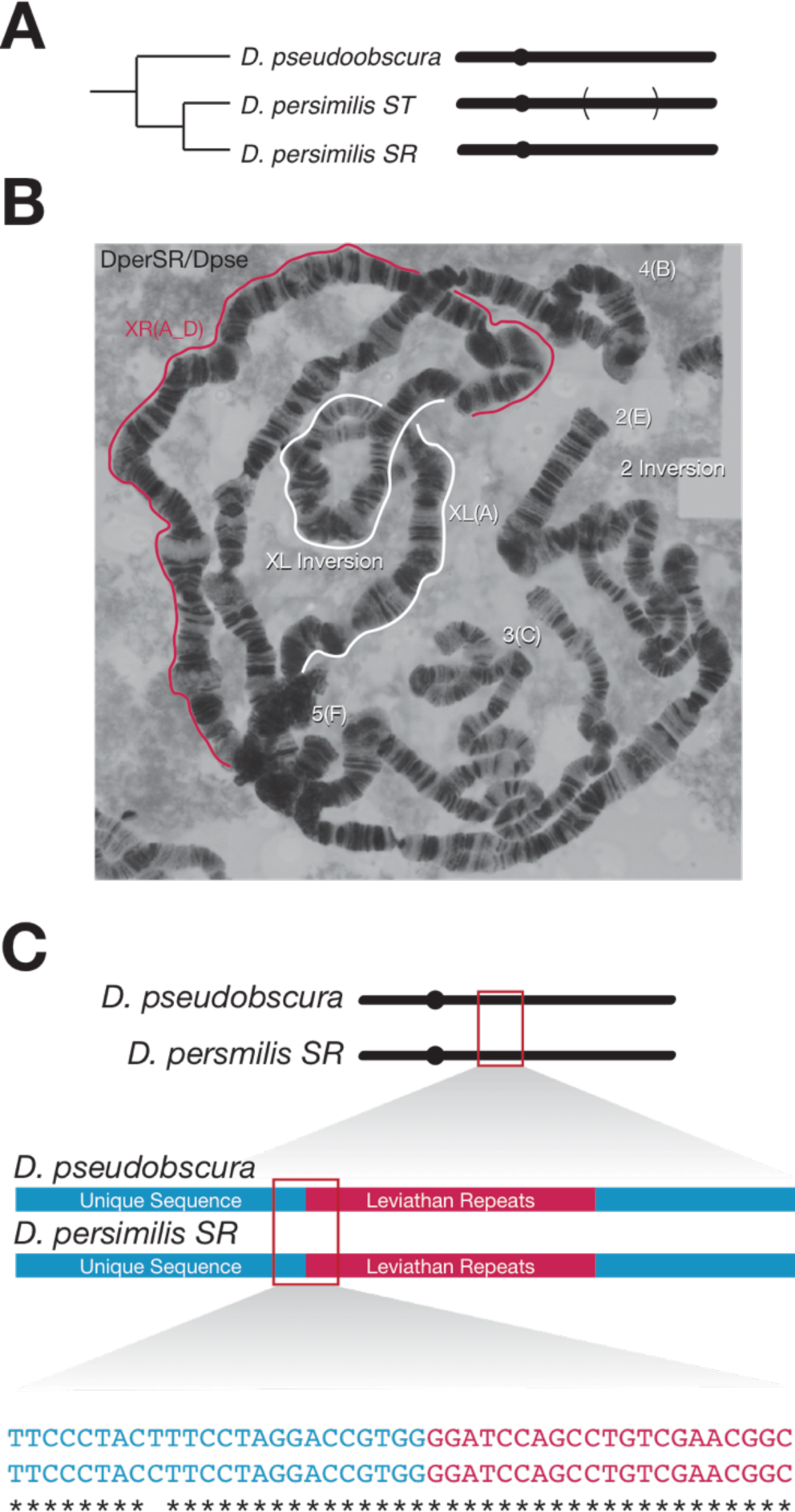
The *D. persimilis Sex-Ratio* (*SR*) chromosome is precisely collinear with *D. pseudoobscura*. *(A)* The right arm of the *X* chromosome (*XR*) of *D. persimilis* is normally inverted as compared to its sister species, *D. pseudoobscura*, but the *D. persimilis Sex-Ratio* chromosome is collinear with its sister species. (B) Polytene chromosome squash of a *D. persimilis SR/D. pseudoobscura* hybrid female demonstrating perfect interspecies collinearity on *XR*. (C) Amplification and sequencing of the proximal breakpoint of the *D. persimilis* inversion reveals that the breakpoints are collinear at the base-pair level.

Here, we resolve the origins of the surprising collinearity of the *D. persimilis SR* chromosome with *D. pseudoobscura XR*. We show that the *D. persimilis SR* chromosome arose, not from a second inversion event, but directly from the ancestrally-arranged chromosome. Surprisingly, we also discovered large blocks of phylogenetic discordance in the regions flanking the *D. persimilis SR* inversion breakpoints, such that they are more closely related to the *D. pseudoobscura*, rather than to the *D. persimilis ST* chromosome. We show that this phylogenetic discordance is not due to the result of gene flow between the two species, but instead arises from incomplete lineage sorting of the derived *D. persimilis ST* inversion from the ancestor of *D. pseudoobscura* and *D. persimilis*. Our results with the *D. persimilis SR* chromosome also show that the recombination-limited regions that flank chromosome inversion breakpoints preserve important clues that allow us to reconstruct the true evolutionary history of these chromosomes, and to more accurately estimate the ages of these chromosomes. We use this insight gleaned from the *D. persimilis SR* analyses to infer the evolutionary histories of the other inversions differences. We show that, contrary to currently the held view, all of the known fixed rearrangements differences between *D. persimilis* and *D. pseudoobscura* arose in the ancestor of the two species long before speciation, but were passed exclusively to *D. persimilis*. Together, our results challenge our current understanding of the evolutionary history of the inversions in *D. persimilis* and *D. pseudoobscura*, and suggest a new model for the role of chromosomal inversions in speciation.

## RESULTS

### A high-resolution examination of polytene chromosomes confirms the apparent collinearity of *D. persimilis Sex-Ratio* with *D. pseudoobscura*

To uncover the evolutionary origins of the *D. persimilis SR* chromosome, we screened for the *Sex-Ratio* trait in wild caught *D. persimilis* flies. We isolated two independent *D. persimilis SR* strains that produce >90% female progeny, and generated high quality mosaic images of polytene chromosomes with squashes of larval salivary glands from them. Consistent with previous reports (18), the *D. persimilis SR* chromosomes in the strains that we isolated differ by one major inversion on *XR* with respect to *D. persimilis ST*, but appear collinear with *D. pseudoobscura* (Figure 1B, Supplementary Figure 1). If *D. persimilis SR* was derived from *D. persimilis ST* through a somewhat imprecise reversion to the ancestral arrangement, the banding patterns of polytene chromosomes in hybrid *D. persimilis SR*/*D. pseudoobscura* females may reveal slight imperfections near the inversion breakpoints. We did not observe any disruption of chromosome pairing near the inversion breakpoints in *D. persimilis SR/D. pseudoobscura* heterozygotes, suggesting that any secondary inversion event may have been in close vicinity of the original breakpoints of the *D. persimilis ST* inversion.

### *D. persimilis Sex-Ratio* and *D. pseudoobscura* are precisely collinear at a single base pair resolution

While our polytene analyses showed no visible aberrations at the breakpoints of the *D. persimilis* inversion, such analyses provide only a coarse view of chromosome structure. Previously, the *D. persimilis ST* inversions breakpoints were mapped at a resolution of 30kb (12). To precisely identify the inversion breakpoints on the *D. persimilis SR* chromosome, we first performed whole genome sequencing of males pooled from two *D. persimilis SR* strains, as well as males pooled from two *D. persimilis ST* strains. Using the approximate genomic coordinates of the inversion breakpoints, we designed multiple primer pairs that span the proximal and distal inversion breakpoint sequences from *D. persimilis SR* and *D. pseudoobscura*. We performed PCR with these primers to successfully amplify single products using *D. persimilis SR* and *D. pseudoobscura* genomic DNA as templates. We were able to amplify sequences corresponding to the proximal breakpoint (Supplementary Figure 2). We identified the precise molecular breakpoints of this inversion by Sanger sequencing the proximal breakpoint PCR products, which revealed the presence of four 319bp *Leviathan* repeats (26) at the breakpoint. More importantly, *D. persimilis SR* and *D. pseudoobscura* sequences that flank the *Leviathan* repeats are precisely collinear to a single base pair resolution (Figure 1C). Although information about the proximal inversion breakpoint also provides accurate information about the position of the distal breakpoint, we were not able to amplify sequences across this region, likely because of the accumulation of repetitive sequences at this breakpoint. Our results from the proximal breakpoint, however, clearly show that a slightly staggered second inversion event is not the basis for the collinearity between the *D. persimilis SR* and *D. pseudoobscura* chromosomes.

### The *D. persimilis Sex-Ratio* chromosome is more closely related to *D. pseudoobscura* than to *D. persimilis* at the inversion breakpoints

If the *D. persimilis SR* inversion originated through a recombination event within *Leviathan* sequences at the inversion breakpoints, such an event can generate the same pattern of perfect collinearity of the flanking sequences. Such repetitive sequences are known to be hotspots for inversion breakpoints (26, 27). (While *Leviathan* repeats are unique to *D. persimilis* and *D. pseudoobscura*, there are more than 850 of these repeats spread across their genomes. Because *XR* alone harbors more than 650 Leviathan repeats spread across the chromosome arm, the probability of a second inversion event on *D. persimilis SR* at the same two *Leviathan* repeats as the original breakpoints appears vanishingly small. However, to directly test whether *D. persimilis SR* is recently derived from *D. persimilis ST* through a secondary inversion event, we inferred phylogenetic relationships in sliding windows across the chromosome, using *D. miranda* as an outgroup (see Methods). As expected, *D. persimilis SR* sequences cluster with those from *D. persimilis ST* across nearly the entire genome (Figure 2A). Surprisingly, we find two large blocks of phylogenetic discordance concentrated at the inversion breakpoints on *XR,* where recombination is expected to be most restricted. In these regions of phylogenetic discordance, *D. persimilis SR* sequences are more closely related to *D. pseudoobscura* rather than to *D. persimilis ST*, with several regions within the inversion also showing the same discordant pattern (Figure 2B). Together with the precise collinearity of *D. persimilis SR* and *D. pseudoobscura,* these results support a single origin of the arrangements of these two chromosomes, followed by limited recombination within the center of the inversion that largely homogenizes the region except for at the breakpoints.

**Figure 2:**
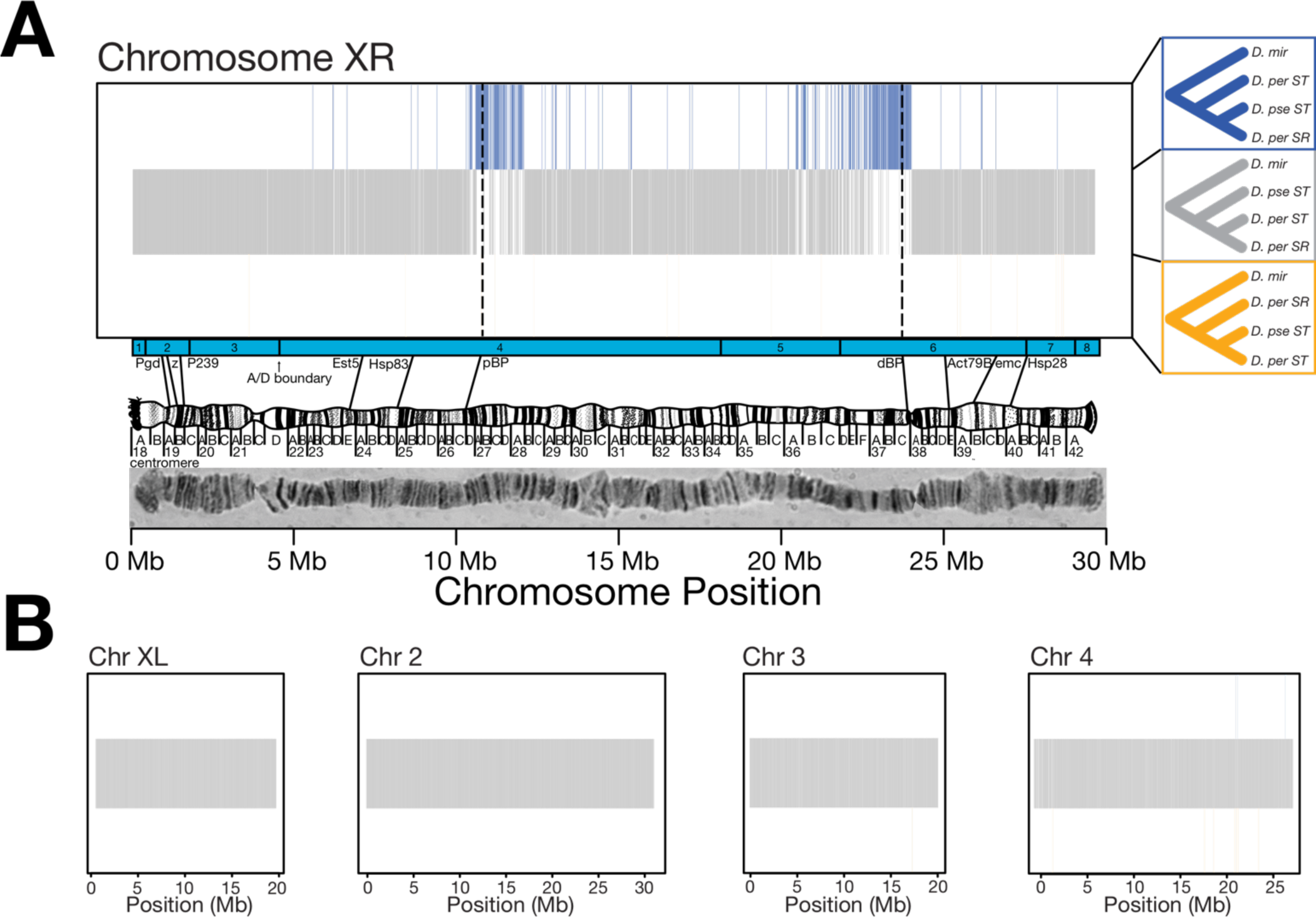
The inversion breakpoints on *XR* show extensive phylogenetic discordance. *(A)* Sliding window phylogeny classification on *XR*. Blue, grey, and orange vertical lines represent the tree topology supported by neighbor-joining trees. Grey trees represent no phylogenetic discordance. Blue trees represent regions where the two collinear chromosomes appear more similar. Large regions centered on the proximal and distal breakpoints (dashed lines) of the *XR* inversion show discordant clustering of *D. persimilis SR* with *D. pseudoobscura* rather than *D. persimilis ST*. *(B)* Large regions of phylogenetic discordance are not observed in the remainder of the genome.

We next asked whether the phylogenetic discordance that we observed with the *D. persimilis SR* chromosome is found anywhere else in the genome. Our sliding window phylogenetic analyses show that there is no significant phylogenetic discordance anywhere else in the genome (Figure 2B). Although these analyses revealed small regions of phylogenetic discordance in other regions of the genome, there is no clustering of consecutive discordant windows, and the discordant windows are not associated with other fixed inversions. *D. persimilis* and *D. pseudoobscura* are known to rarely, but successfully, produce hybrids in nature. A single confirmed F1 hybrid female has been isolated so far, out of tens of thousands of wild caught flies (28). Ongoing interspecies gene flow occurs between collinear chromosomal arrangements that are shared between species but polymorphic within species may, in theory, generate the same pattern of phylogenetic discordance. We were able to test this idea because, like the *Standard* arrangement of *XR*, the *Standard* arrangement on the *3^rd^* chromosome (*3^ST^*) is both shared across *D. persimilis* and *D. pseudoobscura*, and is polymorphic within each species (29). Using *3^ST^* from both species and the *Arrowhead* (*3^AR^*) arrangement of *D. pseudoobscura*, we performed the same phylogenetic analysis across the *3^rd^* chromosome (Supplementary Methods). Sequences at the breakpoints of this shared polymorphic inversion recapitulate the correct species tree, again indicating that the large blocks of phylogenetic discordance at the inversions breakpoints on *XR* are a unique property of the *D. persimilis SR* chromosome (Supplementary Figure 3).

Together with the precisely-shared breakpoints, the relatedness between *D. persimilis SR* and *D. pseudoobscura* at the inversion breakpoints rejects the currently accepted secondary-inversion hypothesis for the origin of the *D. persimilis SR* arrangement, and suggests a single origin for these chromosomes. Our results raise the surprising possibilities that *D. persimilis SR* was derived either through a recent introgression event from *D. pseudoobscura*, or from the common ancestor of *D. persimilis* and *D. pseudoobscura*. Next, we test these two models.

### The *D. persimilis SR* and *ST* arrangements were polymorphic in the ancestor of *D. persimilis* and *D. pseudoobscura*

Because *D. persimilis* and *D. pseudoobscura* can hybridize in nature (28), our results raise the possibility that *D. persimilis SR* originated as a recent introgression of *D. pseudoobscura XR* (Figure 3A). Under the introgression scenario, repeated back-crossing to *D. persimilis* after the initial hybridization event gradually removes *D. pseudoobscura* material through single crossovers outside the inversion, and through double crossovers or gene conversion events inside the inversion. These recombination events homogenize *D. persimilis SR* and *ST*, largely wiping out any hints of a potential cross-species origin of *D. persimilis SR* from *D. pseudoobscura*. However, this history of introgression would be best preserved at the breakpoints of the inversion where suppression of crossovers is greatest (30, 31). The preservation of *D. pseudoobscura* material at the inversion breakpoints would then generate the blocks of phylogenetic discordance we observe on *D. persimilis SR*. The modified *f_d_* statistic is a test to discriminate between introgression versus incomplete lineage sorting (ILS), similar to related “ABBA-BABA” measures, that performs stably when applied in local sequence windows (32). We analyzed our genomic data from *D. persimilis SR* and *ST*, along with *D. pseudoobscura* and *D. miranda* sequences, to estimate the modified *f_d_* across the entire genome (Supplementary Methods). Indeed, we observed significant *f_d_* between *D. pseudoobscura* and *D. persimilis SR* at the same chromosomal inversion breakpoint regions that show phylogenetic discordance (Supplementary Figure 4).

**Figure 3:**
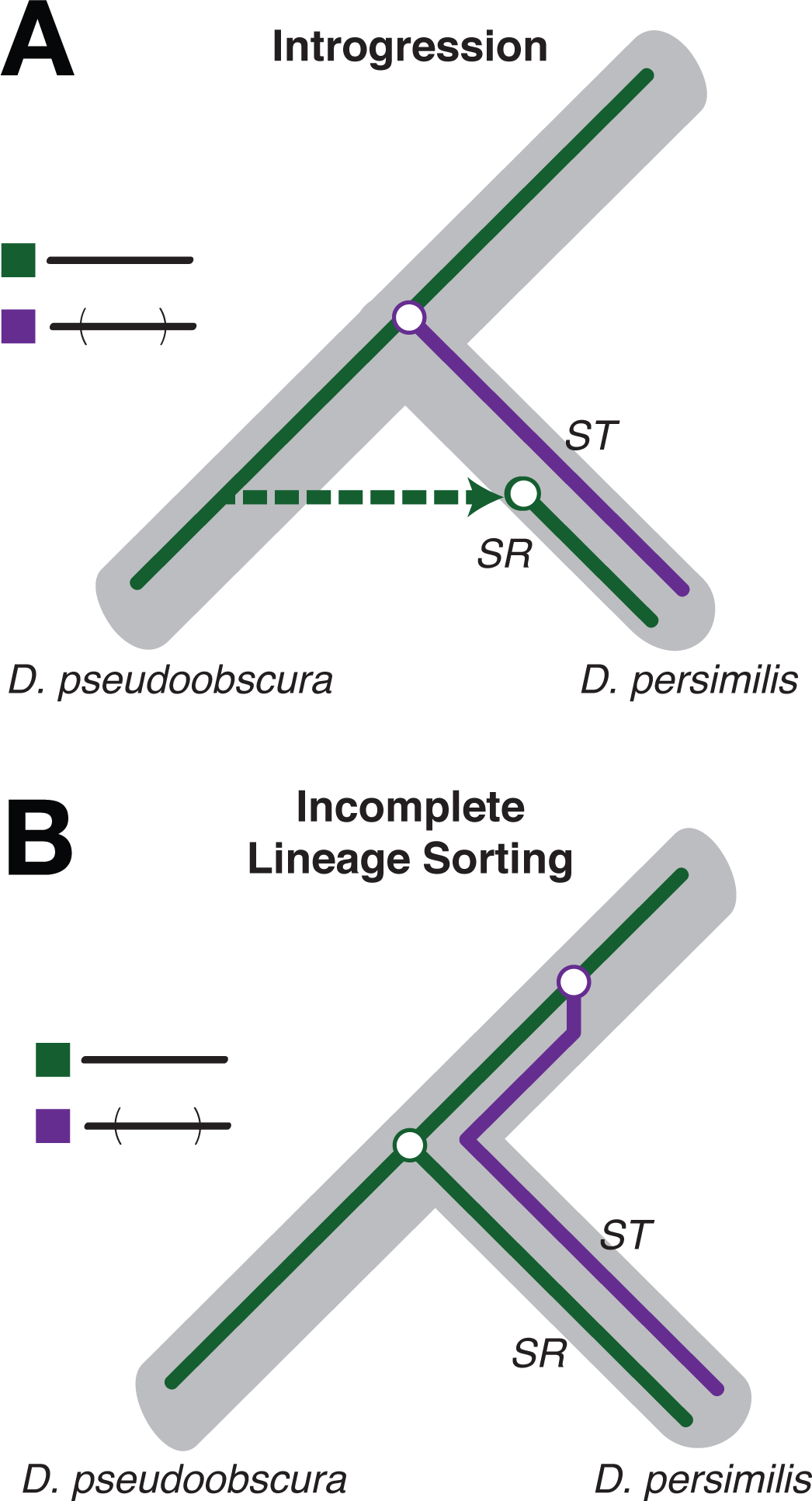
Discordance may be produced by introgression or incomplete lineage sorting of the *XR* arrangements. Under model *(A)*, the *D. persimilis ST* inversion segregates in the ancestral population of the species. Later divergence between *D. persimilis SR* and *D. pseudoobscura* chromosomes and recombination restriction between the two *D. persimilis* chromosomes leads to phylogenetic discordance at the inversion breakpoints. *(B)* An introgression model again predicts discordance if the *D. persimilis SR* chromosome introgressed from *D. pseudoobscura* after species divergence. Recombination between the introgressed chromosome *and D. persimilis ST* will gradually homogenize the two chromosomes excluding the inversion breakpoints.

Interpreting significant values of *f_d_* and related statistics as introgression involves an implicit assumption of free recombination in the ancestral population. However, in regions of limited recombination, such as with inversions, it is incorrect to conclude that introgression is the only cause of such results from these statistical tests. The reason for this is that these regions have a single history, but individual polymorphisms in these regions are summed by the *D* statistic and related statistics in order to assess significance. Because the *D. persimilis SR* chromosome involves a chromosomal inversion that was potentially segregating in the ancestral population, this violates the assumptions required to reliably conclude introgression from such statistics (33). The interpretation of introgression based on any statistic that treats sites as independent may, therefore, be premature and instead may be the result of incomplete lineage sorting (ILS). Indeed, it is not clear if any of the existing statistical approaches can effectively discriminate between introgression and ILS to determine the ancestry of chromosomal inversions or non-recombining chromosomes like the *Y*-chromosome (34).

An alternative explanation that involves the inheritance of the *D. persimilis SR* and *D. pseudoobscura ST* arrangements from the common ancestor of both species can adequately explain the observed patterns. In particular, the phylogenetic discordance that we observe can be explained by the inheritance of the *D. persimilis SR* arrangement from the ancestor of *D. persimilis* and *D. pseudoobscura*, in combination with the loss of one arrangement from *D. pseudoobscura* (Figure 3B). Under this scenario of incomplete lineage sorting in *D. persimilis*, the *ST* inversion originates as a segregating polymorphic chromosome in the ancestral population of *D. persimilis* and *D. pseudoobscura*. The recombination-suppressed regions at the breakpoints of the *D. persimilis ST* inversion begin diverging from the ancestrally arranged chromosomes long before speciation. During this time, the ancestor of *D. persimilis SR* and *D. pseudoobscura ST* chromosomes (which are collinear) continue to freely recombine until the time of speciation, but diverge from the ancestor of the *D. persimilis ST* chromosome. Similar to the introgression scenario, recombination events homogenize *D. persimilis SR* and *ST* after speciation, except at the breakpoints of the inversion, thus leading to the patterns of phylogenetic discordance.

We reasoned that the same recombination-suppressing properties of chromosomal inversions that thwart the application of some statistical approaches may also preserve the information necessary to discriminate between introgression and ILS. In particular, because recombinants at the sequences in the regions near the inversion breakpoints are less frequent, the divergence of the chromosomes can be reliably estimated using these sequences (12). The introgression and ILS hypotheses make distinct and testable predictions about the relative divergence times of each chromosomal arrangement. Under the introgression scenario, we expect the *D. persimilis SR* chromosome to appear much younger than the species divergence time due to a more recent coalescence at introgressed loci. In contrast, the ILS scenario makes two distinct predictions. First, we expect the *D. persimilis SR* chromosome split from the collinear *D. pseudoobscura ST* chromosome much earlier than the species divergence time. Second, we expect the *D. persimilis ST* chromosome and the *D. persimilis SR* chromosome to be more diverged than both the species divergence time *and* the divergence time between *SR* and the *D. pseudoobscura ST* chromosome (Figure 3B).

To test these predictions, we estimated the absolute divergence (*d_xy_*) in 10 kb windows between *D. persimilis* and *D. pseudoobscura* in all collinear regions across the genome and observed a mean *d_xy_* of 2.42x10^−3^ (95% CI: 2.37 − 2.47 ×10^−3^). When standardized to a *D. miranda* divergence set to 2 million years in each window, this corresponds to an allelic divergence time between *D. persimilis* and *D. pseudoobscura* of approximately 500,000 years ago (see Methods). Here, the allelic divergence time represents an upper-bound for the speciation event. We used the sequences flanking the inversion breakpoints (± 250 kb) to estimate *d_xy_* between *D. persimilis SR* and *D. pseudoobscura* and observe a significantly higher (*p*<2.2x10^−16^) mean divergence (4.55 ×10^−3^; 95% CI: 4.22 − 4.89 ×10^−3^) than estimated between species in collinear regions, indicating the *D. persimilis SR* chromosome is older than the speciation time (Table 1). When similarly standardized to the *D. miranda* divergence in each window flanking the breakpoints, we estimate the *D. persimilis SR* chromosome to have diverged ~1.09 million years ago. This is inconsistent with the introgression scenario, and suggests that ILS may better describe the evolutionary origins of the *D. persimilis SR* chromosome. Moreover, the ILS hypothesis makes a second prediction that the *D. persimilis ST* chromosome should be older than the divergence time between the two species. Indeed, we estimate *d_xy_* between *D. persimilis ST* and *D. pseudoobscura* in the same sequences flanking the breakpoint regions (*d_xy_*: 4.94 ×10^−3^; 95% CI: 4.67 − 5.21 ×10^−3^) to be significantly greater (*p*<0.038) corresponding to an older standardized divergence time of ~1.23 million years old.

**Table 1:** Estimates of the relative ages of chromosomal inversions in *D. persimilis* and *D. pseudoobscura* relative to species divergence time. The fixed inversions on the *XL* and *2nd* chromosomes, as well as the polymorphic inversions on *XR* and the Pikes Peak (*3^PP^*) inversion arose before species divergence.

|  | Species divergence | *DperSR* divergence | Chr XR inversion | Chr 2 inversion | Chr XL inversion | Chr 3 PP inversion | Chr 3 AR inversion | Chr 3 AR divergence |
| --- | --- | --- | --- | --- | --- | --- | --- | --- |
| Divergence time (mya) | 0.50 | 1.09 | 1.23 | 1.48 | 1.66 | 0.75 | 0.42 | 0.59 |
| Time prior to species divergence | 0 | 0.59 | 0.73 | 0.98 | 1.16 | 0.25 | -0.08 | 0.09 |

It is important to note three points. First, accurately estimating absolute divergence time in years is known to be fraught with several sources of error and relies on an accurate calibration point in the absence of an estimate of the mutation rate in each species (35). We instead rely on the relative comparison between *d_xy_* estimates, which is sufficient to resolve the questions that we seek to address here. Second, our conclusion that the *D. persimilis ST* inversion existed as a segregating polymorphism in the ancestor of *D. persimilis* and *D. pseudoobscura* is robust to various methods of inferring the divergence between the two species. For example, our results are not significantly different (*p*<0.70) if we use whole genome data or only the collinear regions to estimate the absolute divergence between *D. persimilis* and *D. pseudoobscura* (genome-wide mean *d_xy_*: 2.41 ×10^−3^; 95% CI: 2.36 − 2.46 ×10^−3^). Third, our data only address the order of origins of the various chromosomal inversions, but do not allow us to estimate when the *Sex-Ratio* distortion alleles evolved in these populations. Identifying the causal segregation distortion genes may allow us to address this aspect in the future. Despite these important caveats of our analysis, the two observations of *D. persimilis SR* being older than the species divergence and the *D. persimilis ST* chromosome appearing significantly more diverged than both the collinear regions and the *D. persimilis SR* chromosome reject the introgression scenario and support the ILS explanation.

### All fixed inversions in *D. persimilis* originated as segregating polymorphisms in the ancestral population of *D. persimilis* and *D. pseudoobscura*

Because the *XR* inversion exists only in *D. persimilis* and not in *D. pseudoobscura*, it is often immediately assumed that this inversion must have originated in the *D. persimilis* lineage after speciation (24, 36). The idea that the *XR* inversion on the *Standard* chromosome of *D. persimilis* originated as a segregating polymorphic inversion in the ancestral population prior to speciation goes against this widely-accepted notion. The two other fixed inversions on the *XL* and *2^nd^* chromosomes in *D. persimilis* are thought to be even older than the *XR* inversion (36). We used the same approach that utilizes the sequences flanking inversion breakpoints to also estimate the divergence of the inversions on the *XL* and *2^nd^* chromosomes. Consistent with the idea that the *XL* and *2^nd^* chromosome inversions are older than the *XR* inversion, we observed greater mean levels of *d_xy_* for both fixed inversions (*XL*: 1.08x10^−2^; *2*: 6.82x10^−3^) than for *XR* (*d_xy_*: 4.94 ×10^−3^). Likewise, standardizing to the speciation time with *D. miranda*, we estimate that the inversions on *XL* and the *2^nd^* chromosomes originated approximately 1.66 and 1.48 million years ago, respectively (Table 1; Figure 4). Our results show that all of these fixed inversions originated in the ancestral population long before the speciation event that separated *D. persimilis* and *D. pseudoobscura*. Furthermore, the relative divergence pattern of *XL* > 2 > *XR* that we infer is consistent with the conclusions of previous studies (12, 37).

**Figure 4:**
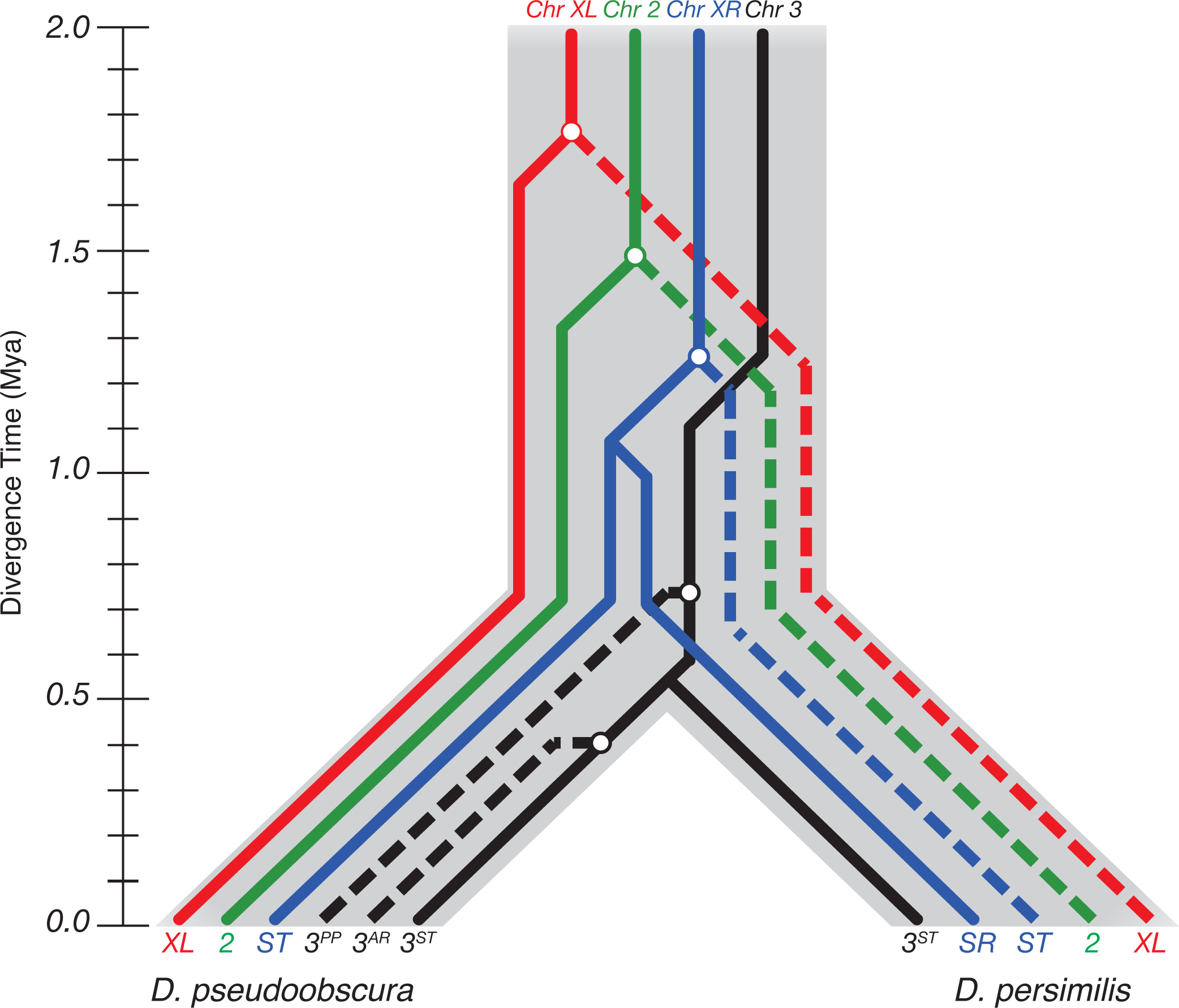
Incomplete lineage sorting of the inversions of *D. persimilis* and *D. pseudoobscura*. The fixed inversions on the *XL* and *2nd* chromosomes, as well as the polymorphic inversions on *XR* and the Pikes Peak (*3^PP^*) inversion arose before species divergence. Incomplete lineage sorting produced the observed inversion patterns in the species present today.

Because it may be argued that genome-wide divergence estimates are a poor proxy for speciation time, we calculated an estimate using another method. *D. persimilis* and *D. pseudoobscura* harbor numerous inversion polymorphisms on the *3^rd^* chromosome that are exclusive to each species, many of which are derived from the *Standard* arrangement (*3^ST^*) that continues to segregate in both *D. persimilis* and *D. pseudoobscura* populations (29, 38). *3^ST^* is the only *3^rd^* chromosome arrangement shared between the two species. Because the *3^ST^* arrangement was present in the ancestral population, and was inherited by both *D. persimilis* and *D. pseudoobscura*, the amount of divergence between the *3^ST^* arrangements of the two species provides another opportunity to determine an upper bound to the estimates of speciation time. We estimated absolute divergence in inversion-associated sequences from the *3^ST^* strains of *D. persimilis* and *D. pseudoobscura*, and observed a mean *d_xy_* of 2.49 x 10^−3^ (95% CI: 2.34 − 2.65 ×10^−3^). Standardizing to the speciation time with *D. miranda* in these regions, we estimate the *3^ST^* inversion between *D. persimilis* and *D. pseudoobscura* to have diverged approximately 590,000 years ago. This estimate of the upper-bound of speciation time is consistent with those using genome-wide sequences, and is far younger than the ages of any of the fixed inversion differences. This difference in the ages of the fixed inversions and the time of speciation is not subtle: while the genome-wide allelic divergence is estimated at around 500,000 years, the *XL*, *XR* and *2^nd^* chromosome inversions are at least twice as old as this estimate (Figure 4). Even if we were to double the estimate of speciation time to about a million years, the fixed chromosomal inversion predate the split by several hundred thousand years. These results suggest that all of the fixed, derived inversions in *D. persimilis* must have freely segregated in the ancestral population for a substantial period of time before speciation.

## DISCUSSION

The study of chromosomal inversions in the classic systems of *D. pseudoobscura* and *D. persimilis* has deeply informed our understanding of the evolutionary forces that shape natural variation, the evolution of new species, and selfish chromosome dynamics. Our results have several important implications for all of these fields. We provide a solution to the strange collinearity of the *D. persimilis SR* and *D. pseudoobscura ST* chromosomes first observed by Dobzhansky (18, 29). We show that this collinearity is a consequence of the direct descent of these chromosomes from one of the ancestrally segregating arrangements, and not due to two independent inversions at the same breakpoints. A similar maintenance of chromosomal arrangements across species resulting from an ancient inversion polymorphism has also been demonstrated in *Anopheles* mosquitos (39). Segregation distorters are often associated with inversions because new inversions that tightly link a segregation distorter gene with existing enhancer alleles enjoy a selective advantage (21). In contrast to most other *Sex-Ratio* systems associated with derived inversions, our results suggest that the *D. persimilis SR* system evolved on the background of an ancestral arrangement. Similarly, recent studies of the *t*-haplotype in *M. musculus* also support an ancient origin of inversions associated with segregation distortion (40). These results indicate that segregation distorters may not only become associated with new inversions, but can arise on existing chromosome inversion polymorphisms.

In addition to clarifying the evolutionary history of *Sex-Ratio* chromosomes in these two species, the divergence we show between fixed arrangement differences in *D. pseudoobscura* and *D. persimilis* suggest a model for the role of chromosomal inversions in the evolution of hybrid incompatibilities. Any model exploring this role must explain at least two empirical patterns: a) the fixed inversions between *D. persimilis* and *D. pseudoobscura* have higher divergence as compared to collinear regions of the genome, and b) most genes that underlie reproductive isolation between *D. persimilis* and *D. pseudoobscura* reside within these inversion differences (9–11). We show that these inversions were freely segregating in the ancestral population long before speciation, and that the genes contributing to reproductive barriers must have evolved within them afterwards.

Here, we propose a simple model to explain the above two empirical patterns (Figure 5). Chromosomal inversions can arise and persist in ancestral populations for long periods of time driven by selection in heterogeneous environments (41). During this period, the genomic regions spanning the inversions and the corresponding regions on the un-inverted chromosomes can accumulate genetic divergence aided by the suppression of recombination in heterozygotes (41–45). Populations with old segregating inversions diverge within inverted regions, but stay genetically similar in collinear regions (42, 45). These chromosomal inversions may undergo incomplete lineage sorting when the ancestral population is split into two allopatric populations (46). At the initial time of separation, the genes within the chromosomal inversions are already highly diverged, whereas the genes within the collinear regions are nearly identical, with little or no divergence. The highly diverged genes associated with chromosomal inversions are fewer mutational steps away from reaching an incompatible state and are, therefore, likely to evolve to an incompatible state more quickly than those in the collinear regions of the genome. This accumulation of hybrid incompatibilities occurs in isolation, unopposed by the selective cost of producing unfit offspring, and in a manner consistent with the Dobzhansky-Muller model (7, 8). The collinear regions will retain their low divergence signature from the ancestral population until speciation is complete, then these regions will begin to fix lineage specific variants. Our simple model is consistent with all empirical results, and is sufficient to explain both patterns. Under our model, the heterogeneity in divergence across the genome caused by ancestrally segregating inversions makes the evolution of alleles that cause reproductive isolation more likely in the regions encompassed by these inversions rather than in the collinear regions of the genome.

**Figure 5:**
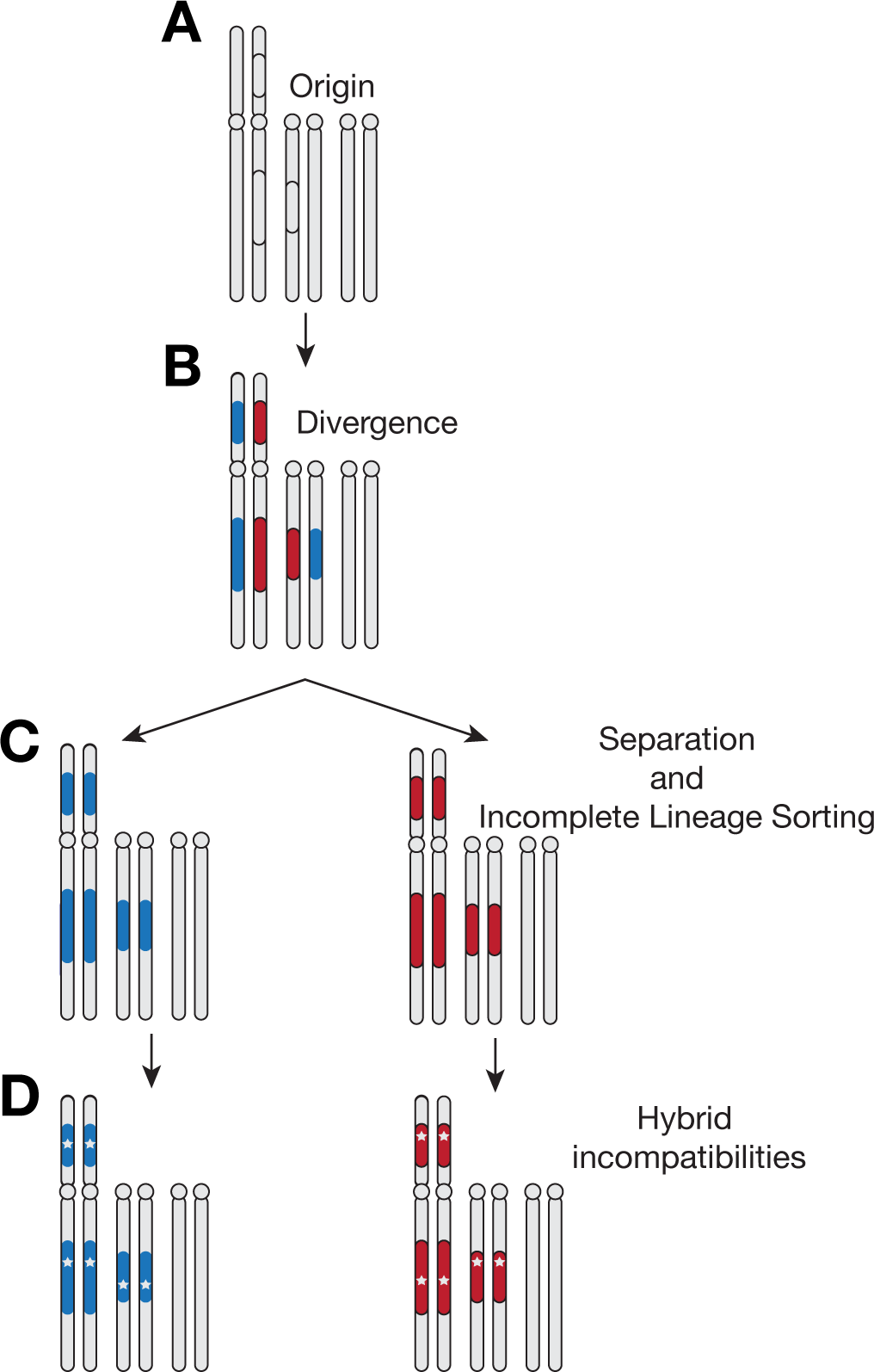
Inversions accelerate the formation of hybrid incompatibilities. *(A)* Polymorphic inversions arise in the ancestor of the two species. *(B)* Restricted recombination between the inversions leads to accumulating divergence (red, blue) distinct from collinear regions of the genome (grey). *(C)* Incomplete sorting of the inversions between two isolated populations generates immediate divergence between the two populations. *(D)* Preexisting divergence increases the chance of hybrid incompatibilities forming in the inverted regions as compared to the collinear regions.

The claim that highly diverged genes are likely to evolve to incompatible state more quickly than those with little divergence rests on the implicit assumption that the evolution of hybrid incompatibilities is a multi-step process that requires multiple changes. Three lines of evidence support the view. First, theory shows that changes at a minimum of two genes are required to produce a hybrid incompatibility, and that it is easier to evolve more complex incompatibilities that involve changes at multiple genes (47). These ideas have strong empirical support (1). For example, the genetic architecture of hybrid sterility between *D. pseudoobscura pseudoodobscura* and *D. pseudoobscura bogotana*–one of the youngest hybridizations to be studied–involves interactions between *at least* six genes (48). Second, nearly all hybrid incompatibility genes that have been identified so far show the rapid accumulation of many amino acid changes, and represent some of the most highly diverged genes in the genome (49, 50). Ultra-fine scale mapping studies that dissect how many of these changes within these genes contribute to hybrid sterility or hybrid inviability have not yet been performed. However, there are no known cases of hybrid incompatibility genes that involve one or only a few amino acid changes. Third, both theory and empirical data show that hybrid incompatibilities accumulate faster than linearly with divergence between populations (51–53). All other things being equal, populations that display higher genomic divergence are, therefore, more likely to have evolved hybrid incompatibilities as compared to those that have little or no genomic divergence (54). Together, these lines of evidence support the idea that the evolution of hybrid incompatibilities is a multi-step process. By accumulating genetic divergence even before the initial population split, the genes associated with ancestrally segregating chromosomal inversions are fewer steps away from reaching an incompatible state. In contrast, genes in collinear regions of the genome show little or no divergence between recently split populations and must start accumulating changes from scratch if they are to eventually an incompatible state.

Both the ‘gene flow during speciation’ and ‘gene flow after speciation’ models rely on gene exchange in collinear regions to account for the higher divergence of fixed inversions, and their association with hybrid incompatibility genes. Multiple studies have presented evidence for *some* level of gene flow between *D. pseudobscura* and *D. persimilis* and rare *F_1_* hybrids have been caught in the wild in locations where these species coexist (12, 28, 37, 55–60). However, the extent to which gene flow occurs and its importance in the reinforcement of reproductive isolation in this species pair have been debated (56, 61, 62). Here, we find little evidence for extensive gene flow or introgression spanning several megabases of sequence that would be required to explain the phylogenetic discordance we observed on the *D. persimilis SR* chromosome (Figure 2). Similarly, strong evidence for gene flow during the accumulation of hybrid incompatibilities as necessitated by the ‘gene flow during speciation’ model is also lacking.

Our model can also account for a third empirical pattern that sympatric species are more likely to harbor fixed chromosomal inversion differences as compared to allopatric species (10, 11, 63). Under our model, when two temporarily isolated populations inherit segregating chromosomal inversions that were likely balanced polymorphisms in the ancestral population, these populations start with highly diverged genomic regions at birth (i.e. associated with inversions) where genes underlying isolating mechanisms may evolve quickly. In contrast, populations that inherit fully collinear genomes have little or no divergence between them at birth and may, therefore, require more time to evolve isolating mechanisms.

If single species often fragment into temporarily isolated populations and merge again, then the populations that inherit ancestrally segregating inversion differences are more likely to survive as separate species even if they later become fully sympatric. In contrast, those with collinear genomes are less likely to evolve hybrid incompatibilities during temporary allopatry and may collapse back into single species on secondary contact. Such cases will not be observed, unless allopatry is maintained long enough to allow the evolution of hybrid incompatibilities. Together, this process is predicted to generate a pattern of sympatric species pairs that are enriched for inversion differences, and allopatric species pairs that have collinear genomes.

Our model also makes a distinct prediction regarding young allopatric species that inherit ancestrally segregating inversions: if such cases are found, hybrid incompatibility genes must be enriched in regions spanned by the inversion differences despite no gene flow between these populations. Because hybrid incompatibilities may accumulate across the genome over time through the snowball effect, this enrichment of hybrid incompatibility genes at inversions may decay over time in older species. This prediction is not expected under the models that rely on gene flow during or after speciation.

The idea that chromosomal inversions are often associated with hybrid incompatibility genes is a widely-held view among evolutionary geneticists (41, 64). There are four lines of reasoning for the widespread acceptance of this notion. First, experimental studies involving genetic mapping of loci that underlie reproductive barriers may be localized to genomic regions that contain fixed chromosomal inversions (11, 65–67). Such studies provide the most direct line of evidence for the association of reproductive isolation genes with chromosomal inversions. Second, genomic regions spanning chromosomal inversions often show signatures of higher divergence or reduced introgression (13, 36, 58). As our results show, this line of evidence may be susceptible to erroneous interpretations when the evolutionary histories and the ages of these inversions are unknown. Third, sympatric species show higher incidence of fixed inversions than allopatric species. While there are limited data supporting such a pattern (68, 69), this line of evidence for the association of hybrid incompatibility genes with chromosomal inversions is indirect and prone to observational biases. Fourth, theoretical studies show that it may be possible for hybrid incompatibility genes to evolve and persist despite gene flow during or after speciation (70, 71). These theoretical results, however, are not a substitute for direct empirical evidence. We, therefore, consider direct genetic mapping studies that localize hybrid incompatibility genes to regions spanning chromosomal inversions as the only reliable line of evidence supporting the association of chromosomal inversions with reproductive isolation genes.

Such genetic studies that map loci that contribute to reproductive isolating barriers, and overlay those loci on the locations of chromosomal inversions are surprisingly rare. To our knowledge, the only direct study of this nature in animal taxa involves the *D. pseudoobscura-D. persimilis* hybridization, where genetic mapping studies have shown that loci that contribute to reproductive isolation are enriched on chromosomes that carry fixed inversion differences between these species (7, 11, 37, 60, 72). In the absence of other such studies, it is not clear whether this pattern is specific to this particular species pair, or is a pattern of a broadly held pattern. We, therefore, find that the amount of evidence for the association of hybrid incompatibility genes with fixed chromosomal inversions is not proportionate to how widely this pattern is believed to be true.

This paucity of mapping studies describing the locations of hybrid incompatibility genes relative to chromosomal inversions is not entirely surprising. A necessary step in understanding the molecular basis of speciation involves the identification of the genes that contribute to reproductive barriers. Most speciation geneticists who aim to identify such genes may either focus on studying species pairs that lack chromosomal inversion differences, or abort such studies when these genes map to chromosomal inversions because there is little hope of precisely identifying the causal genes. Fortunately, uncovering evidence of the association of reproductive isolation genes with chromosomal inversions requires neither the precise identification of the genes nor determining the precise breakpoints of chromosomal inversions. Coarse mapping of quantitative trait loci that underlie reproductive isolation across several species pairs, and overlaying these loci with the approximate locations of chromosomal inversion differences between these species may prove sufficient to establish the generality of this pattern (65).

In summary, we propose that higher divergence and hybrid incompatibilities are emergent properties of chromosomal inversions when they are segregating in ancestral populations and inherited through incomplete lineage sorting. Our model explains previously observed empirical patterns without invoking gene flow across populations during or after speciation and forces a reconsideration of the role of inversions in speciation, perhaps not as protectors of existing hybrid incompatibility alleles, but as fertile grounds for their formation.

## MATERIALS AND METHODS

### Isolation and maintenance of Sex-Ratio chromosome strains

Wild caught *D. persimilis* strains were provided as a generous gift by Dean Castillo, collected in the Sierra Nevada mountain range and near Mt. St. Helena, CA. We tested individuals from these strains for the presence of *Sex-Ratio* chromosomes by crossing males to standard *D. persimilis* females. We isolated two individual *D. persimilis Sex-Ratio* strains and generated stable stocks through eight to twelve generations of inbreeding. All stocks were raised on standard cornmeal media at 18 degrees C.

### Polytene chromosome analyses

We used two crosses of *D. persimilis SR*/*ST* heterozygotes to compare the *D. persimilis SR* chromosome with *D. pseudoobscura* and *D. persimilis ST* chromosomes. In the first cross, a *D. persimilis SR*/*ST sepia* (*se*) heterozygous female was crossed to a *D. pseudoobscura ST se* male. Of the two *XL*/*XR* karyotypes possible from this cross, we examined females heterozygous for *XL* and homozygous for *XR* inversions. These females allow us to evaluate whether the *D. persimilis SR* and *D. pseudoobscura ST* chromosomes are homosequential. In a second cross, a *D. persimilis SR*/*ST se* heterozygous female was crossed to a *D. persimilis ST se* male. Of the two *XL*/*XR* karyotypes possible from this cross, we examined females homozygous for *XL* and heterozygous for *XR* inversions. These females allow us to examine the *D. persimilis SR* and *D. persimilis ST* heterozygotes. We prepared salivary squashes from larvae from these two crosses using standard techniques, with modifications described by Harshman (1977) and Ballard and Bedo (1991) (61–63).

### DNA extraction and sequencing

To generate whole genome shotgun sequencing libraries for *D. persimilis* strains, we pooled one male each from two *SR* strains and two *ST* strains (from Sierra Nevada and Mt St Helena collections). We extracted DNA from these flies using the 5 Prime Archive Pure DNA extraction kit according to the manufacturer’s protocol (ThermoFisher, Waltham, MA). All libraries were generated with the Illumina TruSeq Nano kit (Epicentre, Illumina Inc, CA) using the manufacturers protocol, and sequenced as 500bp paired end reads on an Illumina HiSeq 2000 instrument.

### Sequence alignment and SNP identification

Low-quality bases were removed from the ends of the raw paired end reads contained in FASTQ files using *seqtk* (https://github.com/lh3/seqtk) with an error threshold of 0.05. Illumina adapter sequences and polyA tails were trimmed from the reads using Trimmomatic version 0.30 (64). The read quality was then manually inspected using FastQC. Following initial preprocessing and quality control, the reads from each pool were aligned to the *D. pseudoobscura* reference genome (v 3.2) using *bwa* version 0.7.8 with default parameters (65). Genome wide, the average fold coverage was ~180x and ~133x for the *D. persimilis ST* and *SR* pools, respectively (Supplementary Table 1). For reads mapping to X chromosome scaffolds, the average fold coverage was ~97x and ~74x for *D. persimilis ST* and *SR*, respectively (Supplement Table 2).

After the binary alignments were sorted and indexed with SAMtools (66), single nucleotide polymorphisms (SNPs) were called using *freebayes* (v. 0.9.21; (67) with the expected pairwise nucleotide diversity parameter set to 0.01, based on a previous genome-wide estimate from *D. pseudoobscura* (40). The samples were modeled as discrete genotypes across pools by using the “–J” option and the ploidy was set separately for *X* chromosome scaffolds (1*N*) and autosomes (2*N*). SNPs with a genotype quality score less than 30 were filtered from the dataset. We restricted all downstream analyses to sites that had coverage greater than *1N* and less than 3 standard deviations away from the genome wide mean for all samples (Table S1). Across the genome we identified a total of 3,598,524 polymorphic sites, 703,908 and 844,043 of which were located on chromosomes *XR* and *XL*, respectively.

The *D. pseudoobscura* reference assembly does not contain complete sequences for either of the arms of the *X* or *4^th^* chromosomes. Instead, each is composed of a series of scaffold groups that differ both in size and orientation relative to one another (68). Schaeffer et al. (2008) previously determined the approximate locations and ordering of each of these scaffolds (68). We used their map to convert the scaffold-specific coordinates of each site to the appropriate location on the corresponding chromosome to construct a continuous sequence.

### Estimating the phylogenetic relationship of Sex-Ratio chromosomes

We estimated the genetic distance between each pairwise grouping in 10 kb windows using Nei’s *D_A_* distance, which has been shown to accurately recover the topology of phylogenetic trees from allele frequency data (69, 70). To root the tree with an outgroup, we aligned publically available short reads of *D. miranda* (SRX965461; strain SP138) to the *D. pseudoobscura* reference genome. In each window, we constructed neighbor-joining trees (71) using distance matrices constructed from the estimated genetic distances (*D_A_*) and classified the phylogeny based on the topology it supported. If a window contained fewer than 10 segregating sites, we did not construct a tree or estimate the genetic distance. For each tree we performed 10,000 bootstrap replicates and only included those windows with a support value of 0.75 or higher.

### Divergence Estimates

We estimated absolute divergence with Nei’s *d_xy_*, a measure of the average number of pairwise nucleotide substitutions per site (72, 73). *d_xy_* was measured between each population grouping in 10 Kb, nonoverlapping windows across the genome. To convert estimates of absolute divergence into divergence times, the *d_xy_* values were scaled to a 2 My species split between *D. pseudoosbcura* and *D. miranda* in each window. Methods for estimating introgression with the modified *f_d_* statistic are found in the Supplementary Methods.

### Identification and verification of inversion breakpoints

The proximal and distal breakpoints have both been characterized previously, and the regions in *D. pseudoobscura* contain unique sequence flanking a series of 302-bp repeats known as Leviathan repeats, present throughout the genomes of both *D. pseudoobscura* and *D. persimilis*. We designed primers to capture both the array of repeats as well as portions of unique sequence. We extracted DNA from all three genotypes and amplified the proximal breakpoint region using primers designed to anneal to the *D. pseudoobscura* genomic sequence flanking the Leviathan repeats (F5’-GATCTAATCCAGAAAGTTCGCTTGCG-3’, R5’-AGTGTGACCCATTTTAAGCGG-3’). These primers amplified a single, approximately 1500bp, product in *D. pseudoobscura* and *D. persimilis SR*, but not *D. persimilis ST*. PCR products were Sanger sequenced using the forward and reverse PCR primers at the DNA Sequencing Core Facility, University of Utah. The reads were aligned both to one another and to sequence from the *D. pseudoobscura* genome assembly around the proximal breakpoint. The sequenced PCR product was confirmed to contain both the repeats and sections of the unique sequence flanking the repeat region at the proximal breakpoint.

## ACKNOWLEDGEMENTS

This work was supported by the National Institutes of Health (Genetics Training Grant 5T32GM007464-40 (CJL), R01 GM115914 (NP), R01 GM 098478 (SWS), a Mario Capecchi endowed assistant professorship (NP), and the Pew Biomedical Scholars Program. We thank Dean Castillo for generously providing wild-caught *D. persimilis* flies. We are particularly grateful to Molly Schumer, and Matt Hahn for his third reviewer services (@3rdreviewer) and for originally asking us to consider an incomplete lineage sorting hypothesis.

## REFERENCES

1. Coyne JA, Orr HA (2004) Speciation (Sinauer).

2. Dobzhansky T (1937) Genetics and the Origin of Species (Columbia University Press).

3. White MJD (1978) Modes of speciation (W.H. Freeman, San Francisco).

4. Dobzhansky T (1933) On the sterility of the interracial hybrids in *Drosophila pseudoobscura*. Proc Natl Acad Sci USA 19(4):397–403.

5. Stebbins GL (1970) Variation and Evolution in Plants: Progress During the Past Twenty Years. Essays in Evolution and Genetics in Honor of Theodosius Dobzhansky (Springer, Boston, MA), pp 173–208.

6. Stebbins GL (1958) The inviability, weakness, and sterility of interspecific hybrids. Adv Genet 9:147–215.

7. Dobzhansky T (1936) Studies on Hybrid Sterility. II. Localization of sterility factors in *Drosophila pseudoobscura* hybrids. Genetics 21(2):113–135.

8. Muller HJ (1942) Isolating mechanisms, evolution and temperature. Biological Symposia, The Jaques Catell Press. (Lancaster, PA), pp 71–125.

9. Wu CI, Beckenbach AT (1983) Evidence for Extensive Genetic Differentiation between the Sex-Ratio and the Standard Arrangement of *Drosophila pseudoobscura* and *D. persimilis* and Identification of Hybrid Sterility Factors. Genetics 105(1):71–86.

10. Brown KM, Burk LM, Henagan LM, Noor MAF (2004) A test of the chromosomal rearrangement model of speciation in *Drosophila pseudoobscura*. Evolution 58(8):1856–1860.

11. Noor MAF, Grams KL, Bertucci LA, Reiland J (2001) Chromosomal inversions and the reproductive isolation of species. PNAS 98(21):12084–12088.

12. Machado CA, Haselkorn TS, Noor MAF (2007) Evaluation of the Genomic Extent of Effects of Fixed Inversion Differences on Intraspecific Variation and Interspecific Gene Flow in *Drosophila pseudoobscura* and *D. persimilis*. Genetics 175(3):1289–1306.

13. Kulathinal RJ, Stevison LS, Noor MAF (2009) The Genomics of Speciation in Drosophila: Diversity, Divergence, and Introgression Estimated Using Low-Coverage Genome Sequencing. PLOS Genetics 5(7):e1000550.

14. Machado CA, Kliman RM, Markert JA, Hey J (2002) Inferring the history of speciation from multilocus DNA sequence data: the case of *Drosophila pseudoobscura* and close relatives. Mol Biol Evol 19(4):472–488.

15. Ayala FJ, Coluzzi M (2005) Chromosome speciation: humans, Drosophila, and mosquitoes. Proc Natl Acad Sci USA 102 Suppl 1:6535–6542.

16. Rieseberg LH (2001) Chromosomal rearrangements and speciation. Trends Ecol Evol (Amst) 16(7):351–358.

17. Navarro A, Barton NH (2003) Chromosomal speciation and molecular divergence--accelerated evolution in rearranged chromosomes. Science 300(5617):321–324.

18. Sturtevant AH, Dobzhansky T (1936) Geographical Distribution and Cytology of “Sex Ratio” in *Drosophila pseudoobscura* and Related Species. Genetics 21(4):473–490.

19. Policansky D, Ellison J (1970) “Sex ratio” in *Drosophila pseudoobscura*: spermiogenic failure. Science 169(3948):888–889.

20. Bastide H, Gérard PR, Ogereau D, Cazemajor M, Montchamp-Moreau C (2013) Local dynamics of a fast-evolving sex-ratio system in *Drosophila simulans*. Mol Ecol 22(21):5352–5367.

21. Jaenike J (2001) Sex Chromosome Meiotic Drive. Annual Review of Ecology and Systematics 32(1):25–49.

22. Hamilton WD (1967) Extraordinary sex ratios. A sex-ratio theory for sex linkage and inbreeding has new implications in cytogenetics and entomology. Science 156(3774):477–488.

23. Presgraves DC, Gérard PR, Cherukuri A, Lyttle TW (2009) Large-Scale Selective Sweep among Segregation Distorter Chromosomes in African Populations of *Drosophila melanogaster*. PLOS Genetics 5(5):e1000463.

24. Babcock CS, Anderson WW (1996) Molecular evolution of the Sex-Ratio inversion complex in *Drosophila pseudoobscura*: analysis of the Esterase-5 gene region. Mol Biol Evol 13(2):297–308.

25. Kovacevic M, Schaeffer SW (2000) Molecular population genetics of X-linked genes in *Drosophila pseudoobscura*. Genetics 156(1):155–172.

26. Garfield DA, Noor MA (2007) Characterization of novel repetitive element Leviathan in *Drosophila pseudoobscura*. Drosophila Information Service 90:1–9.

27. Aguado C, et al. (2014) Validation and Genotyping of Multiple Human Polymorphic Inversions Mediated by Inverted Repeats Reveals a High Degree of Recurrence. PLOS Genetics 10(3):e1004208.

28. Dobzhansky T (1973) Is there Gene Exchange between Drosophila pseudoobsura and Drosophila persimilis in Their Natural Habitats? The American Naturalist 107(954):312–314.

29. Dobzhansky T (1944) Chromosomal races in Drosophila pseudoobscura and Drosophila persimilis (Washington Publ., Carnegie Inst.).

30. Navarro A, Barbadilla A, Ruiz A (2000) Effect of Inversion Polymorphism on the Neutral Nucleotide Variability of Linked Chromosomal Regions in Drosophila. Genetics 155(2):685–698.

31. Navarro A, Betrán E, Barbadilla A, Ruiz A (1997) Recombination and Gene Flux Caused by Gene Conversion and Crossing Over in Inversion Heterokaryotypes. Genetics 146(2):695–709.

32. Durand EY, Patterson N, Reich D, Slatkin M (2011) Testing for Ancient Admixture between Closely Related Populations. Mol Biol Evol 28(8):2239–2252.

33. Martin SH, Davey JW, Jiggins CD (2015) Evaluating the Use of ABBA–BABA Statistics to Locate Introgressed Loci. Mol Biol Evol 32(1):244–257.

34. Hall AB, et al. (2016) Radical remodeling of the Y chromosome in a recent radiation of malaria mosquitoes. PNAS 113(15):E2114–E2123.

35. Graur D, Martin W (2004) Reading the entrails of chickens: molecular timescales of evolution and the illusion of precision. Trends Genet 20(2):80–86.

36. Noor MAF, Garfield DA, Schaeffer SW, Machado CA (2007) Divergence Between the *Drosophila pseudoobscura* and *D. persimilis* Genome Sequences in Relation to Chromosomal Inversions. Genetics 177(3):1417–1428.

37. McGaugh SE, Noor MAF (2012) Genomic impacts of chromosomal inversions in parapatric Drosophila species. Philos Trans R Soc Lond B Biol Sci 367(1587):422–429.

38. Moore BC, Taylor CE (1986) Drosophila of Southern California: III. Gene arrangements of *Drosophila persimilis*. J Hered 77(5):313–323.

39. Fontaine MC, et al. (2015) Extensive introgression in a malaria vector species complex revealed by phylogenomics. Science 347(6217):1258524.

40. Kelemen RK, Vicoso B (2017) Complex History and Differentiation Patterns of the t-Haplotype, a Mouse Meiotic Driver. Genetics:genetics.300513.2017.

41. Kirkpatrick M, Barton N (2006) Chromosome Inversions, Local Adaptation and Speciation. Genetics 173(1):419–434.

42. Fuller ZL, Haynes GD, Richards S, Schaeffer SW (2016) Genomics of Natural Populations: How Differentially Expressed Genes Shape the Evolution of Chromosomal Inversions in *Drosophila pseudoobscura*. Genetics:genetics.116.191429.

43. Fuller ZL, et al. (2014) Evidence for Stabilizing Selection on Codon Usage in Chromosomal Rearrangements of Drosophila pseudoobscura. G3:g3.114.014860.

44. Schaeffer SW, et al. (2003) Evolutionary genomics of inversions in *Drosophila pseudoobscura*: Evidence for epistasis. Proc Natl Acad Sci U S A 100(14):8319–8324.

45. Fuller ZL, Haynes GD, Richards S, Schaeffer SW (2017) Genomics of Natural Populations: Evolutionary Forces that Establish and Maintain Gene Arrangements in *Drosophila pseudoobscura*. Mol Ecol:n/a-n/a.

46. Guerrero RF, Hahn MW Speciation as a Sieve for Ancestral Polymorphism. Mol Ecol:n/a-n/a.

47. Orr HA (1995) The population genetics of speciation: the evolution of hybrid incompatibilities. Genetics 139(4):1805–1813.

48. Phadnis N (2011) Genetic Architecture of Male Sterility and Segregation Distortion in *Drosophila pseudoobscura* Bogota– USA Hybrids. Genetics 189(3):1001–1009.

49. Presgraves DC (2010) The molecular evolutionary basis of species formation. Nat Rev Genet 11(3):175–180.

50. Maheshwari S, Barbash DA (2011) The genetics of hybrid incompatibilities. Annu Rev Genet 45:331–355.

51. Matute DR, Butler IA, Turissini DA, Coyne JA (2010) A test of the snowball theory for the rate of evolution of hybrid incompatibilities. Science 329(5998):1518–1521.

52. Moyle LC, Nakazato T (2010) Hybrid incompatibility “snowballs” between Solanum species. Science 329(5998):1521–1523.

53. Wang RJ, White MA, Payseur BA (2015) The Pace of Hybrid Incompatibility Evolution in House Mice. Genetics 201(1):229–242.

54. Roux C, et al. (2016) Shedding Light on the Grey Zone of Speciation along a Continuum of Genomic Divergence. PLoS Biol 14(12):e2000234.

55. Wang RL, Wakeley J, Hey J (1997) Gene flow and natural selection in the origin of *Drosophila pseudoobscura* and close relatives. Genetics 147(3):1091–1106.

56. Noor MA, Johnson NA, Hey J (2000) Gene flow between *Drosophila pseudoobscura* and *D. persimilis*. Evolution 54(6):2174–2175-2177.

57. Powell JR (1983) Interspecific cytoplasmic gene flow in the absence of nuclear gene flow: evidence from Drosophila. Proc Natl Acad Sci USA 80(2):492–495.

58. Stevison LS, Hoehn KB, Noor MAF (2011) Effects of Inversions on Within- and Between-Species Recombination and Divergence. Genome Biol Evol 3:830–841.

59. Noor MA (1995) Speciation driven by natural selection in Drosophila. Nature 375(6533):674–675.

60. Noor MA, et al. (2001) The genetics of reproductive isolation and the potential for gene exchange between *Drosophila pseudoobscura* and *D. persimilis* via backcross hybrid males. Evolution 55(3):512–521.

61. Kulathinal RJ, Singh RS (2000) Reinforcement with gene flow? a reply. Evolution 54(6):2176–2177.

62. Kulathinal RJ, Singh RS (2000) A Biogeographic Genetic Approach for Testing the Role of Reinforcement: The Case of *Drosophila pseudoobscura* and *D. persimilis*. Evolution 54(1):210–217.

63. Hooper DM, Price TD (2017) Chromosomal inversion differences correlate with range overlap in passerine birds. Nature Ecology & Evolution 1(10):1526.

64. Hoffmann AA, Rieseberg LH (2008) Revisiting the Impact of Inversions in Evolution: From Population Genetic Markers to Drivers of Adaptive Shifts and Speciation? Annual Review of Ecology, Evolution, and Systematics 39(1):21–42.

65. Fishman L, Stathos A, Beardsley PM, Williams CF, Hill JP (2013) Chromosomal rearrangements and the genetics of reproductive barriers in mimulus (monkey flowers). Evolution 67(9):2547–2560.

66. Lowry DB, Willis JH (2010) A Widespread Chromosomal Inversion Polymorphism Contributes to a Major Life-History Transition, Local Adaptation, and Reproductive Isolation. PLOS Biology 8(9):e1000500.

67. McDermott SR, Noor MAF (2012) Mapping of within-species segregation distortion in *D. persimilis* and hybrid sterility between *D. persimilis* and *D. pseudoobscura*. J Evol Biol 25(10):2023–2032.

68. Castiglia R (2014) Sympatric sister species in rodents are more chromosomally differentiated than allopatric ones: implications for the role of chromosomal rearrangements in speciation. Mammal Review 44(1):1–4.

69. Davey JW, et al. (2017) No evidence for maintenance of a sympatric Heliconius species barrier by chromosomal inversions. Evolution Letters 1(3):138–154.

70. Feder JL, Nosil P (2009) Chromosomal Inversions and Species Differences: When Are Genes Affecting Adaptive Divergence and Reproductive Isolation Expected to Reside Within Inversions? Evolution 63(12):3061–3075.

71. Pinho C, Hey J (2010) Divergence with Gene Flow: Models and Data. Annual Review of Ecology, Evolution, and Systematics 41(1):215–230.

72. Orr HA (1987) Genetics of Male and Female Sterility in Hybrids of *Drosophila pseudoobscura* and *D. persimilis*. Genetics 116(4):555–563.

73. Painter TS (1934) A New Method for the Study of Chromosome Aberrations and the Plotting of Chromosome Maps in Drosophila Melanogaster. Genetics 19(3):175–188.

74. Harshman LG (1977) A technique for the preparation of Drosophila salivary gland chromosomes. Drosophila Information Service 52(164).

75. Ballard JWO, Bedo DG (1991) Population cytogenetics of Austrosimulium bancrofti (Diptera: Simuliidae) in eastern Australia. Genome 34(3):338–353.

76. Bolger AM, Lohse M, Usadel B (2014) Trimmomatic: A flexible trimmer for Illumina Sequence Data. Bioinformatics:btu170.

77. Li H, Durbin R (2009) Fast and accurate short read alignment with Burrows-Wheeler transform. Bioinformatics 25(14):1754–1760.

78. Li H, et al. (2009) The Sequence Alignment/Map format and SAMtools. Bioinformatics 25(16):2078–2079.

79. Garrison E, Marth G (2012) Haplotype-based variant detection from short-read sequencing. arXiv:12073907 [q-bio]. Available at: http://arxiv.org/abs/1207.3907 [Accessed September 5, 2017].

80. Schaeffer SW, et al. (2008) Polytene Chromosomal Maps of 11 Drosophila Species: The Order of Genomic Scaffolds Inferred From Genetic and Physical Maps. Genetics 179(3):1601–1655.

81. Nei M, Tajima F, Tateno Y (1983) Accuracy of estimated phylogenetic trees from molecular data. II. Gene frequency data. J Mol Evol 19(2):153–170.

82. Kalinowski ST (2002) Evolutionary and statistical properties of three genetic distances. Mol Ecol 11(8):1263–1273.

83. Saitou N, Nei M (1987) The neighbor-joining method: a new method for reconstructing phylogenetic trees. Mol Biol Evol 4(4):406–425.

84. Nei M (1987) Molecular Evolutionary Genetics (Columbia University Press).

85. Nei M, Li WH (1979) Mathematical model for studying genetic variation in terms of restriction endonucleases. PNAS 76(10):5269–5273.

